# Methylation guide RNA evolution in archaea: structure, function, and genomic organization of 110 C/D box sRNA families across six *Pyrobaculum* species

**DOI:** 10.1101/121921

**Authors:** Lauren M Lui, Andrew V. Uzilov, David L. Bernick, Andrea Corredor, Todd M. Lowe, Patrick P. Dennis

**Affiliations:** Department of Biomolecular Engineering, University of California Santa Cruz, Santa Cruz, CA, 95064, USA; Department of Biology, Whitman College, Walla Walla, WA, 99362, USA

**Keywords:** C/D box sRNA, archaea, RNA-seq, comparative genomics, computational search model

## Abstract

Archaeal homologs of eukaryotic C/D box small nucleolar RNAs (C/D box sRNAs) guide precise 2′-*O*-methyl modification of ribosomal and transfer RNAs. Although C/D box sRNA genes constitute one of the largest RNA gene families in archaeal thermophiles, most genomes have incomplete sRNA gene annotation because reliable, fully automated detection methods are not available. We expanded and curated a comprehensive gene set across six species of the crenarchaeal genus *Pyrobaculum*, particularly rich in C/D box sRNA genes. Using high-throughput small RNA sequencing, specialized computational searches, and comparative genomics, we analyzed 526 *Pyrobaculum* C/D box sRNAs, organizing them into 110 families based on synteny and conservation of guide sequences which determine methylation targets. We examined gene duplications and rearrangements, including one family that has expanded in a pattern similar to retrotransposed repetitive elements in eukaryotes. New training data and inclusion of kink-turn secondary structural features enabled creation of an improved search model. Our analyses provide the most comprehensive, dynamic view of C/D box sRNA evolutionary history within a genus, in terms of modification function, feature plasticity, and gene mobility.

## INTRODUCTION

In eukaryotic cells, ribosome assembly occurs in the nucleolus, a specialized structure located within the nucleus. At this site, ribosomal RNA (rRNA) is transcribed, modified, processed, folded, and assembled along with ribosomal proteins into the large and small ribosomal subunits. The nucleolus also contains a large number of small RNAs (snoRNAs) that are required for the modification and maturation of rRNA, and implicated as chaperones in folding (reviewed in (1)). These snoRNAs are incorporated into dynamic ribonucleoprotein (RNP) complexes that act as molecular processing machines along the ribosome assembly line. Most snoRNAs contain guide sequences that base pair with rRNA, facilitating precise modification of ribonucleotides within the region of complementarity. The snoRNAs divide into two classes: C/D box snoRNAs, which guide 2ʹ-*O*-methylation of ribose, and H/ACA box snoRNAs, which guide the conversion of uridine to pseudouridine (2, 3). Although archaeal cells do not contain an organized nucleolar structure, they possess and utilize both C/D box and H/ACA box sno-like RNAs (sRNAs) in the modification of rRNA and assembly of ribosomal subunits (reviewed in (1, 4)). Notably, bacteria do not use RNA-guided modification, so archaea have become the most convenient, minimally complex models for studying this innovation in enzymatic flexibility.

Archaeal C/D box sRNAs are generally about 50 nucleotides (nts) in length and contain highly conserved C (RUGAUGA consensus) and D (CUGA consensus) box sequences at the 5ʹ and 3ʹ ends of the molecule, and less conserved versions (designated Cʹ and Dʹ) near the center of the molecule (5). These RNAs fold into a hairpin as a result of the formation of a kink-turn (K-turn) structural motif through the interaction of the C and D box sequences and a K-loop motif through the interaction of the Dʹ and Cʹ box sequences (Figure 1A). The K-turn and the K-loop are each recognized by the protein L7Ae (6). The binding of L7Ae stabilizes the RNA structure and allows two copies each of Nop56/58 (also called Nop5) and fibrillarin to bind, completing the assembly of the active ribonucleoprotein (RNP) complex (7). The fibrillarin protein is an *S*-adenosyl methionine dependent RNA methylase, and is responsible for the catalytic activity of the RNP complex.

**Figure 1:**
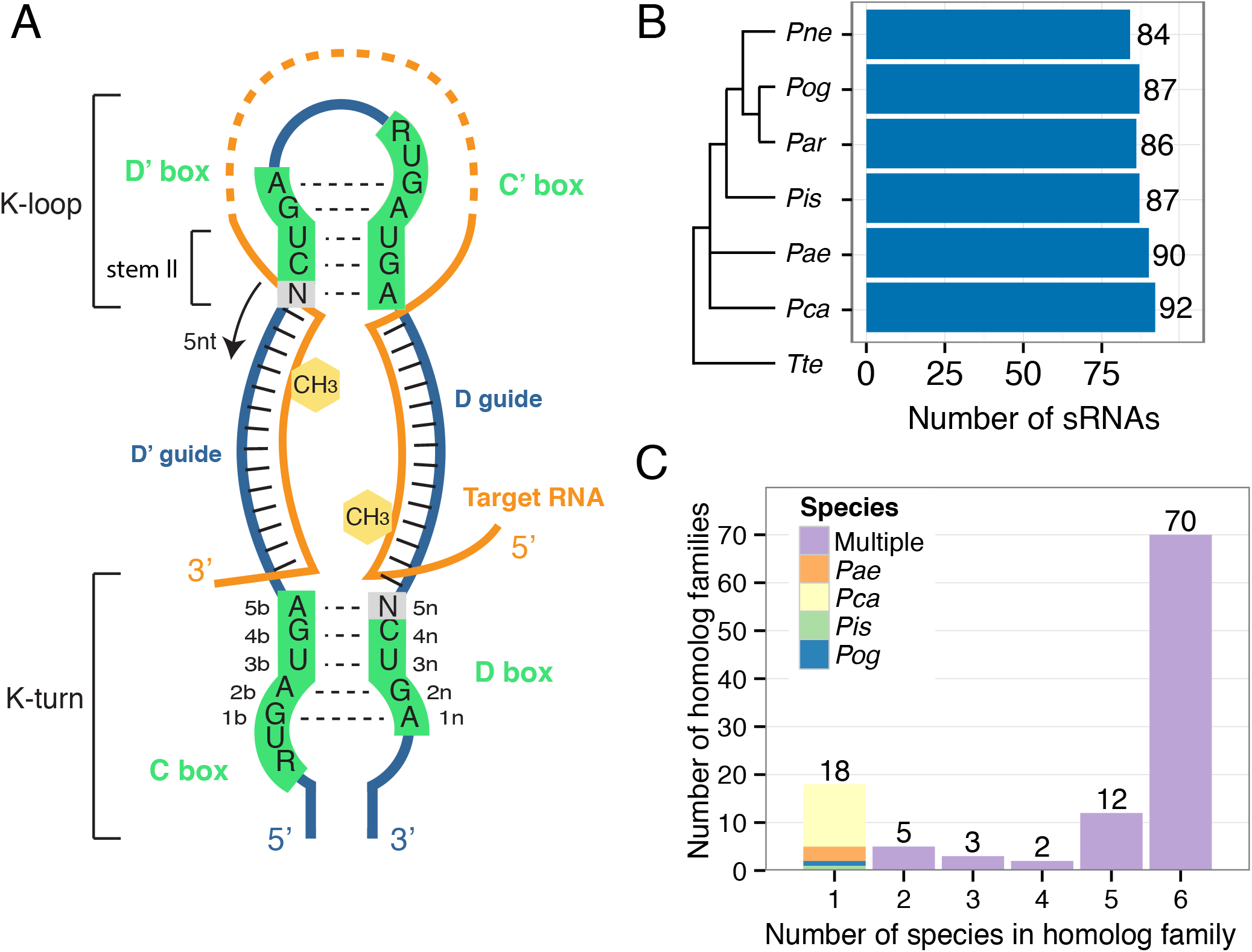
Organization of *Pyrobaculum* C/D box RNAs into 110 homologous families. (A) The typical structure of an archaeal C/D box sRNA is depicted. The structure contains two K-motifs, the K-turn formed by the interaction between the C and D box sequences and the K-loop formed by the interaction between the Dʹ and Cʹ motifs (black dashed lines). The two guide regions located respectively between the Dʹ and C boxes and between the D and Cʹ boxes (green), base pair with the target RNA (orange) and methylation (yellow hexagon) occurs in the target nucleotide that base pairs with the guide five nts upstream from the start of the Dʹ or D box sequence. This is the “N+five” rule. (B) The number of identified sRNAs in each of the six species of *Pyrobaculum* is indicated, and closely mirrors the adjacent species tree, as determined by 16S rRNA alignment (51). *Thermoproteus tenax* (*Tte*) is included as an outgroup. (C) C/D box sRNAs were organized into 110 homologous families based on sequence similarity of the guides and predicted targets in rRNA and tRNAs. C/D box sRNA numbers indicate to which family each belongs. Thus, *Pae* sR01, *Par* sR01, etc. belong to the sR01 family. C/D box sRNAs were first grouped into families using the original annotation numbering in *Pae* (1-65) (23). All other C/D box sRNAs were grouped into families starting at number 100. The majority of sRNAs fall into families with representatives on all six species.

The two guide regions between the C and Dʹ boxes and between the Cʹ and D boxes are unstructured and each is available to base pair with an approximately 8–12nt long target sequence (Figure 1A). In addition to rRNA targets, a significant proportion of archaeal sRNAs have guide regions that are complementary to transfer RNA (tRNA) (7, 8). Methyl modification in the target RNA occurs at the nucleotide position that base pairs with the guide five nucleotides upstream from the first base of the Dʹ or D box sequence. This is known as the “N+five” rule and methylation targets are referred to as the D and Dʹ targets (2). Many C/D box sRNAs with canonical box features have guide regions that lack complementarity to rRNA and tRNA sequences; these are known as “orphan” guides and may target other RNAs, but to date no conserved targets to other RNAs have been identified within the Archaea (1).

The evolution of C/D box sRNAs affects ribosome function and the host genome. In all domains of life, ribose methylations help stabilize RNA structure and are most often found in functionally important regions of the ribosome, such as the peptidyl transferase center in domain V of the large ribosomal subunit (9). Although elimination of individual 2′-*O*-methylations by C/D box sRNA deletions appear to have little effect on the cell, global dysregulation of the methyltransferase fibrillarin has profound effects, possibly including cancer in humans (10, 11). In addition to their role in ribose methylation, the propagation of C/D box sRNA genes may have a profound impact on the evolution and architecture of the genome. In mammals and nematodes, some C/D box sRNAs appear to duplicate via a retrotransposon-like mechanism (12–14). These duplications may lead to new functions of the sRNAs, and like other transposons, play a role in genome evolution (15).

Although other studies have detected C/D box sRNA gene duplication and instances of their overlap with protein-coding genes (16–18), none have been comprehensive with the intent of understanding sRNA evolution and function within a genus. The *Pyrobaculum* genus is diverse yet provides an ideal genetic distance where orthology and synteny of sRNA genes can still be clearly established. We supported C/D box sRNA gene predictions with a combination of comparative genomics and small RNA sequencing (RNA-seq) data from five *Pyrobaculum* species (18, 19): *P. aerophilum* (*Pae*), *P. arsentaticum* (*Par*), *P. calidifontis* (*Pca*), *P. islandicum* (*Pis*), and *P. oguniense* (*Pog*). In addition, the species *P. neutrophilum* (*Pne*; formerly *Thermoproteus neutrophilus* (20)) was used to supplement comparative genomics analyses. After identifying a reference set of 526 high-confidence C/D box sRNA genes, the most comprehensive set in any archaeal genus to date (16, 21, 22), we aligned and curated them into homologous families based on sequence similarity and methylation target prediction. Finally, we used this extensive data set to make a range of new observations regarding sRNA evolution between the six *Pyrobaculum* genomes based on (i) variation in sequence features within homologous sRNA families, (ii) inferred conservation of function in terms of rRNA modification and ribosome assembly, and (iii) sRNA gene context and plasticity (mobility, duplication, flanking gene orientation). This new reference set also enabled us to create and test a new, highly sensitive computational search model for archaeal C/D box sRNAs.

## MATERIALS AND METHODS

### Computational prediction and organization of C/D box sRNA homolog families

To generate a complete or nearly complete set of sRNA gene predictions within the genus *Pyrobaculum* (Supplementary Table S1), we used computational covariance models and small RNA sequencing data of *Pae*, *Par*, *Pca*, *Pis*, and *Pog* (data reported in (18) except for *Pog*). In a previous study where we used small RNA-seq data from four species of *Pyrobaculum*, we identified several unannotated transcripts that were likely to be conserved C/D box sRNAs with box features (specifically the K-turn motif formed by box base pairing) that were divergent from the canonical C/D box model (18). Therefore, we developed a covariance model that accommodates orphan guides and incorporates box sequences, K-turn and K-loop structure, and length of spacers (guides and variable loop) (Figure 1A). The two G:A pairs of the K-turn and K-loop, were annotated in the structural alignment used as input for the model. The variable spacer regions of archaeal C/D box sRNAs, the guides and variable loop, were annotated to be any base. The length of these spacer regions in the model is based on the longest observed length in the training set.

The initial covariance model was created by using a hand-curated multiple structural alignment of the 62 *Pae* C/D box RNAs reported in the genome sequencing paper and subsequent computational analysis (16, 23). These served as input to `cmbuild` from the Infernal v1.0 and v1.1 software packages (24, 25) with the hand-curated option `--rf` or `–-hand` specified, respectively. The covariance model was calibrated with `cmcalibrate`. A final covariance model was built from a complete set of *Pae* sRNAs found from examining sequencing data and using comparative genomics with the other *Pyrobaculum* species (Supplementary File S1). This high-quality training set only contains C/D box RNAs that are either conserved within the *Pyrobaculum* genus or have confirming small RNA-seq data. We used `cmsearch` to scan the genomes and small RNA sequencing data using the glocal (`-g`) and no HMM filter (`–-nohmm`) options.

We used Infernal v1.0 (24) to pick up between 0-5 more predictions per species since it is slightly more sensitive than Infernal v1.1 (25) (Supplementary Table S2). It is possible to increase the sensitivity of Infernal v1.1 using the `--max` option, but the number of candidates to evaluate manually becomes prohibitive as the number of false positives can increase into the hundreds or thousands. Infernal 1.0 and 1.1 produce different subsets of candidates; we used both for the final prediction set and manually curated the sRNAs that were predicted by only one or the other.

To improve specificity, candidates from genome scans that overlapped other annotated non-coding RNAs or overlapped Genbank RefSeq protein-coding genes by more than 80% were discarded, unless they were supported by syntenic ortholog predictions in other *Pyrobaculum* genomes. The model predicts sRNA box features; these were manually checked and adjusted when required.

To find additional orthologs of sRNAs genes within the *Pyrobaculum* genomes not found by the covariance model, the genomes were searched using sRNA sequences as queries to BLASTN (26). Genome Genbank/INSDC numbers are AE009441.1 (*Pae*), CP000660.1 (*Par*), CP000561.1 (*Pca*), CP000504.1 (*Pis*), CP001014.1 (*Pne*), and CP003316.1 (*Pog*). Top hits were manually curated, based on predicted promoters, conservation, and sequencing evidence. Families of C/D box sRNA homologs were created based on sequence similarity of guide sequences and predicted target sites of modification. The first 62 families were assigned numbers (1-65) based on the previously reported *Pae* sRNA numbering (18). *Pae* sR10 was renamed *Pae* sR57a to reflect homology to *Pae* sR57 and the sR57 family. *P*ae sR41 was removed from annotations because of a lack of sequencing reads and no recognizable Cʹ or Dʹ boxes. *Pae* sR55 was also removed from further analysis because it does not have a recognizable D box and has no predicted orthologs in the six other *Pyrobaculum* genomes in this study. Additional families containing newly identified sRNAs were numbered 100 to 141.

The sequences, family organization, and genomic location of these sRNAs can be found in Supplementary Table S1, at the Lowe Lab Archaeal snoRNA-like C/D box RNA Database (<u>http://lowelab.ucsc.edu/snoRNAdb/</u>) and (27). UCSC Archaeal Genome Browser tracks enabling the visualization of the annotated small RNAs can also be found on the website (28).

### Prediction of methylation targets

To identify the putative sites of 2ʹ-*O*-methylation guided by *Pyrobaculum* C/D box sRNAs, we scanned mature rRNA and tRNA sequences for regions of complementarity to the D and Dʹ guides of the sRNAs. Mature rRNA sequences were obtained by a global alignment of the six *Pyrobaculum* rRNAs and removal of predicted introns. Intron sites are indicated in Supplementary Figures S1 and S2. A uniform numbering system for sites of rRNA methylation was obtained by constructing an alignment of the 16S and 23S rRNA sequences shared across all six species (Supplementary Figures S1 and S2). The predicted locations of modification were mapped on the alignment and assigned a position based on the *Pae* rRNA numbering. For a prediction to be considered credible, generally a minimum complementarity of nine continuous Watson-Crick base pairs centering at or near the “N+five” position was required. The criteria were relaxed in two specific instances. First, if the majority of members in an sRNA group met the prediction criteria, the prediction was extended to minority members that nearly met the criteria (for example, matches containing a mismatch or G:U base pair). Second, it has been noted that many sRNAs use their two guide regions to direct methylations to closely spaced nucleotides within the target RNA (16). Presumably, this enhances target identification and creates greater stabilization of the guide-target interaction within the RNP complex. Consequently, when one guide exhibits strong complementarity to the target, the criterion for the second guide match is relaxed if (i) it is within 100 nucleotides of the first complementarity, (ii) the weaker complementarity contains no more than one mismatch, and (iii) the combined bit score for the two complementarities was 32 or higher (where a Watson-Crick base pair is 2, a G:U base pair is 1 and a mismatch is -2). This empirically derived rule-based method is applied with the Python program *findAntisense.py*. Description of the program and related files can be found at https://github.com/lmlui/findAntisense.

### Northern Blot of polycistronic sRNAs

Northern blots were prepared as described in (18). The following DNA oligomers (Integrated DNA Technologies, Inc., Coralville, IA) were used as probes:

*Pae* sR21 sense (5’-GCCAGTGTCCGAAAATTGACGAGCTCACCCTTTGC)
*Pae* sR21 antisense (5’-GCAAAGGGTGAGCTCGTCAATTTTCGGACACTGGC).

### Small RNA sequencing and read processing of *Pyrobaculum* species

Small RNA sequencing data for *Pae*, *Par*, *Pca*, and *Pis* were mined from a previous study from our lab (18). The libraries for these four species were sequenced on the Roche/454 GS FLX sequencer. Small RNA libraries for *Pog* were sequenced by the UC Davis Sequencing Facility on Illumina HiSeq 2000 to produce 2x75 nt paired-end sequencing reads. Sample preparation of small RNA libraries for *Pog* are described in (18, 29). Briefly, the small RNA size fraction was isolated by running total RNA in denaturing gel electrophoresis and extracting the region below tRNAs. Reads with barcodes and linkers removed were mapped to genomes using BLAT (30). The resulting PSL file was processed to determine paired reads. Supporting small RNA-seq data can be interactively visualized using the UCSC Archaeal Genome Browser and a companion track hub (instructions provided at <u>http://lowelab.ucsc.edu/snoRNAdb/TrackHubs.html)</u>.

### Data access

The small RNA-seq data and BigWig files used to create the track hubs discussed in this publication have been deposited in NCBI's Gene Expression Omnibus (31, 32) and are accessible through GEO Series accession number GSE106228 (https://www.ncbi.nlm.nih.gov/geo/query/acc.cgi?acc=GSE106228).

## RESULTS

### Identification and Curation of *Pyrobaculum* C/D box sRNA genes

We identified 526 C/D box sRNA genes from six species of *Pyrobaculum* using evidence from (i) RNA-seq data from *Pae*, *Par*, *Pca*, *Pis* and *Pog*, (ii) an improved computational covariance prediction model, and (iii) comparative genomics. Nearly all of the sRNAs from the five genomes with RNA-seq data (436/442, 99%) are represented by reads in the RNA-seq libraries, and most were conserved in other *Pyrobaculum* species. The new covariance model developed in this study had two advantages over prior search methods: (1) incorporation of K-turn and K-loop structural information, and (2) it does not rely on guide sequence target detection to identify candidates, allowing detection of sRNAs where both guides lack complementarity to rRNA or tRNA sequences (1, 33).

#### Most Pyrobaculum C/D box sRNA homolog families have members in all six species

The 526 *Pyrobaculum* sRNA genes were organized into 110 different homologous families based on sequence conservation of their guide regions and predicted targets of methylation in tRNA and rRNA (Supplementary Table S1). We found 26 previously undetected sRNAs in this study (two in *Pae*, three in *Par*, four in *Pca*, three in *Pis*, four in *Pog*, and ten in *Pne*); the total number of detected C/D box sRNA genes in individual species ranges between 84 and 92 (Figure 1B). Grouping the sRNAs into families allowed us to track the evolution of C/D box sRNA genes (gain, loss, methylation target changes) within the genus and to predict target sites of methylation within rRNA and tRNA with greater certainty.

Most of the homologous families are conserved, with 70 of the 110 families (64%) having representative sRNAs encoded in each of the six *Pyrobaculum* genomes. The 40 remaining families have representatives missing from one or more of the six genomes (Figure 1C). Eighteen of the families are unique with the representative present in only a single species. Each of these 18 singleton sRNAs have small RNA-seq reads and 15 have at least one predicted target to rRNA or tRNA. Within a family, it is common for both guide regions to exhibit a high degree of sequence similarity indicative of a common ancestry. For example, of the 70 families that have representative sRNAs in all six species, 62 exhibit a recognizable degree of sequence similarity in both the D and the Dʹ guide regions among all members, whereas the remaining eight families have a conserved sequence across all species in only one of the two guide regions (see Supplementary Table S1). Even when a particular guide region is conserved, it is frequently punctuated by very short (1-3 nucleotides) insertions, deletions, or substitutions, primarily at the 5ʹ or 3ʹ end of the guides that are not predicted to form part of the guide-target base pairing interaction. Accordingly, not a single guide region is perfectly conserved in any of the 70 families with representatives in all six species, emphasizing that these species have diverged enough to observe differences in selective pressure between functional and non-functional segments of sRNAs.

#### New computational model based on curated Pyrobaculum C/D box sRNAs

One of the important aspects of our detailed curation of archaeal sRNAs was the ability to generate a gold-standard set for training a computational model. After training the model with a curated alignment of *Pae* sRNA genes (see Methods) and obtaining score distributions of true positives and false positives, we determined a high confidence threshold of 17 bits (best specificity), and a moderate confidence threshold of 13 bits (high sensitivity with some loss of specificity) for archaeal C/D box sRNA predictions (Supplementary Table S2 and Supplementary Figure S3).

To test how well this model works on divergent archaea, we used it to search three species in the euryarchaeal genus *Pyrococcus*: *P. abyssi* (*Pab*), *P.furiosus* (*Pfu*), and *P. horikoshii* (*Pho*). The genus contains many sRNA gene predictions and has been a model for studying C/D box sRNA structure and function (21, 22). We found seven C/D box sRNAs among these species (two in *Pab*, four in *Pfu*, and one in *Pho*, Supplementary Table S3). These predictions are conserved with other *Pyrococcus* sRNAs or have small RNA sequencing evidence (Supplementary Figure S4). We noted that on average the *Pyrococcus* sRNAs have bitscores that are 2 points higher than the *Pyrobaculum* sRNAs, which is surprising since the model was trained on *Pyrobaculum* sequencing (Supplementary Figure S4A-D). Upon further analysis, we found that *Pyrococcus* C/D box sRNAs score higher than *Pyrobaculum* sRNAs because they have less variation in their box sequences and more canonical K-turns compared to the *Pyrobaculum* (Figure 2). The larger sequence variation in the *Pyrobaculum*, however, may make this model more sensitive for scanning other Archaea.

**Figure 2:**
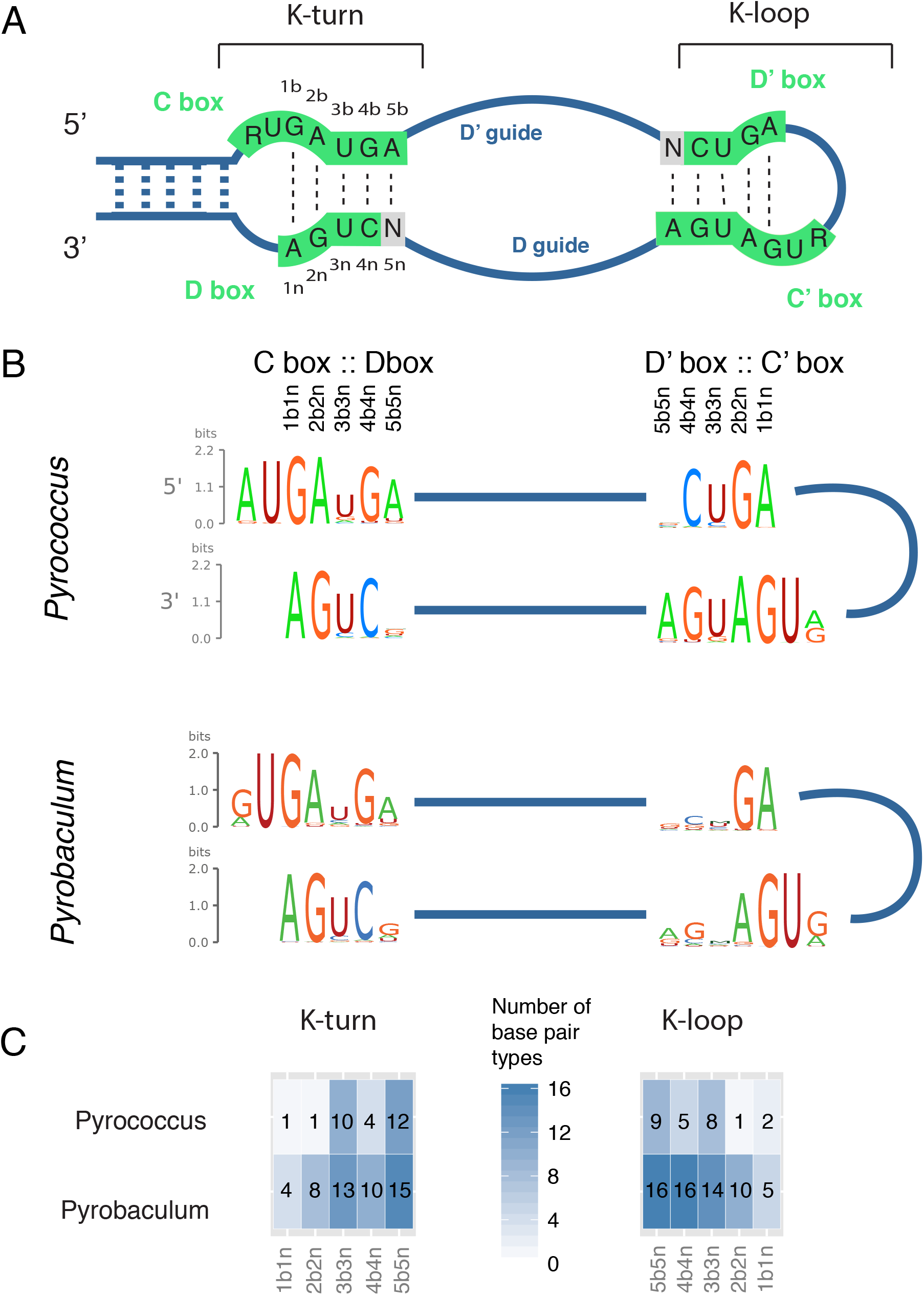
Comparison of *Pyrobaculum* and *Pyrococcus* C/D box sRNA K-turns and K-loops. (A) Diagram of K-turn and K-loop positions in an archaeal C/D box sRNA. In K-turn studies, the base pairs are numbered starting with the G:A base pairs and the first letter in the pair is from the C box (52, 53). The strand with the bulge and the C box is referred to as ‘b,’ and the non-bulged strand with the D box is referred to as ‘n.’ Thus the first two positions are typically position 1b1n being G:A and position 2b2n being A:G. (B) Sequence logos of the box features of the *Pyrobaculum* and *Pyrococcus* genera. Created with RILogo (54). (C) Representation of the variability of base pair type (16 possible) found at each position of the K-turn or K-loop. Darker boxes indicate that position is more variable.

Approximately 3-11% of known C/D box sRNA genes in the *Pyrobaculum* and *Pyrococcus* genomes are not predicted with this covariance model. We examined the false negatives and found that in most cases one or more box features were unusual, often resulting in non-canonical K-motifs (see Supplementary Figure S3 for discussion). To capture these unusual C/D box sRNAs, another model may be needed.

### Prediction and Analysis of C/D box sRNA methylation targets

Methylation targets in rRNA and tRNA were predicted using the “N+five” rule (2) (Figure 1A), and hits were ranked based on extended complementarity between guide sequences and target RNAs (Supplementary Tables S4 and S5). Using criteria described in Materials and Methods, we were able to predict at least one methylation target for 89% (468/526) of the sRNAs. Across all possible guide regions (two per sRNA), 56% (587/1052) were predicted to mediate rRNA methylation, and 16% (173/1052) were predicted to guide tRNA methylation. Using this large set of genes and predicted target methylation sites, we describe results yielding evolutionary and functional insights below.

#### Analysis of targets in ribosomal RNA support the role of sRNAs as rRNA folding chaperones

Positions of predicted methyl modification were mapped onto the 16S and 23S rRNA secondary structure in order to visualize clustering patterns (Supplementary Figures S5 and S6). As noted for other species, methylation sites cluster within functionally important regions, such as the peptidyl transferase center and helix 69 of 23S rRNA. Comparisons with positions of predicted modification in species outside of *Pyrobaculum* indicate that the precise sites of modification are, with a few notable exceptions, generally not conserved, although the clustering pattern is conserved (17).

Notably, approximately 45% of sRNAs (237/526) use their D and Dʹ guides to target sites that are within 100 nts of each other in the primary rRNA sequence (Supplementary Figures S1, S2, S5, S6, and Supplementary Tables S4 and S5). These observations further support prior studies that have suggested dual guide interactions may play an important role in mediating the folding and stabilization of nascent rRNAs and their assembly into ribosomal subunits (17, 34), particularly in thermophiles. A computational study simulating C/D box sRNA chaperone function in rRNA folding also suggests that double guide sRNAs may be especially important for proper long-range interactions in rRNAs (35).

Three sRNAs (sR2, sR53, sR56), conserved in all six species, have D and Dʹ guides that have complementarities and predicted methylation targets that are more than 100 nts apart in the primary rRNA sequence but are close in the secondary structure (Supplementary Figure S7). These long-range interactions occur in the 16S translational fidelity and central core regions, which are important for the function of the ribosome (see Supplementary Figure S7 for more discussion). We suspect that these long-range interactions also play an important role in the tertiary folding of rRNA during the assembly process.

#### *One-third of* Pyrobaculum *C/D box sRNAs target tRNAs*

It has been shown previously that archaeal C/D box RNAs can target modification to tRNAs as well as rRNAs (23). The tRNA methylation targets are at structurally conserved positions that are modified by protein-only tRNA methylases in other organisms (16, 36). In our collection of 110 *Pyrobaculum* sRNA families, 32 are predicted to target modification to 23 different positions in various tRNAs (Supplementary Tables S4 and S5). Of these tRNA-targeting sRNAs, about 80% (115/144) have one guide that targets a tRNA and the other guide has no target. Dual-guide sRNAs with potential to target different sites in the same tRNA were rare (sR46, sR61, sR123, sR126; Supplementary Table S4 and S5), in contrast to the many dual-guide sRNAs targeting rRNA.

The number of different tRNAs that could be targeted by a particular sRNA guide varies over a wide range and reflects the fact that some sequences in tRNAs are unique whereas others are shared among many different tRNA isoacceptors. The region surrounding position 34, the wobble base in the anticodon, is an example of a fairly unique target sequence. Guides from four different sRNAs target position C34 or U34, and each has only a single tRNA target (sR26:C34Trp, sR27:U34Gln, sR45:C34Val, sR46:U34Thr and sR51:C24Glu). Other guides have multiple potential tRNA targets; for example, the D guide of *Pae* sR64 exhibits complementarity to a conserved sequence in sixteen different tRNA families and directs modification to position G51 in the TΨC stem.

#### Instances of mismatched base pairs at the “N+five” position of methylation

*In vivo* and *in vitro* studies have demonstrated that a Watson-Crick base pair at the “N+five” position in the guide-target base pairing region is essential for methylation of the target RNA(7, 37, 38). Nonetheless, a mismatch at the site of methylation within a conserved guide-target complementarity implies that the interaction may be beneficial but that the modification is either not needed or harmful to the function of the target RNA. Studies in yeast that use cross-linking to detect RNA-RNA interactions support this hypothesis (39, 40).

In four instances in the *Pyrobaculum* sRNA set, we find a conserved mismatch at the “N+five” methylation position (sR116:U109 and sR25:C1368 in 16S; sR56:G764, sR33:C2045 in 23S; Supplementary Tables S4 and S5). For example, in the sR33 family, the D and D′ guides target closely spaced positions in 23S rRNA in all six *Pyrobaculum* species (Figure 3). The Dʹ guide of all members is predicted to be incapable of methylation at C2045 because of a C:U mismatch. The Dʹ guide-target interaction is credible because of its strong conservation and its close proximity to the D guide interaction.

**Figure 3:**
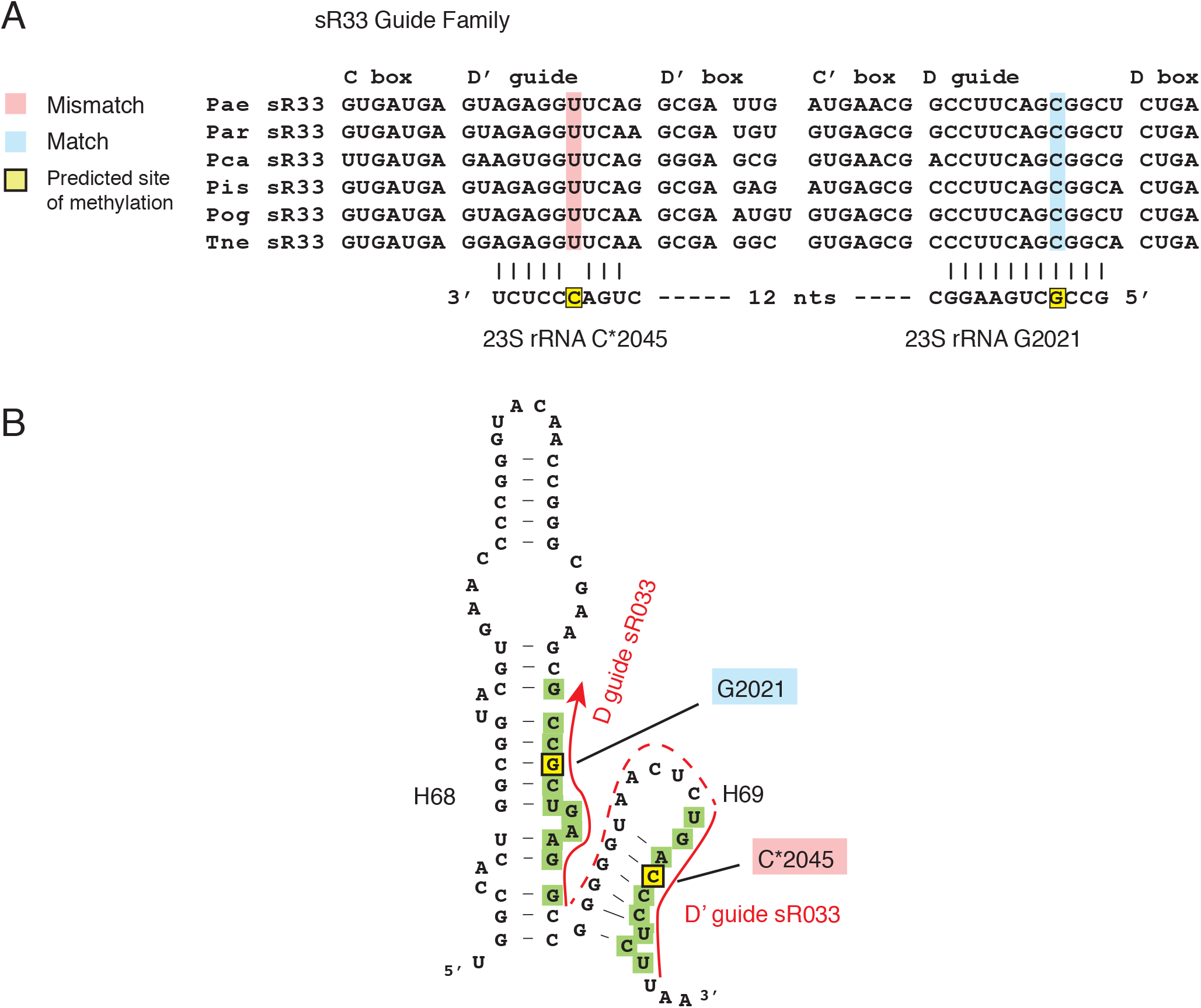
Conserved Instances of mismatch at the site of modification (“N+5” position) indicate that in some cases the target interaction, rather than the modification, is important. (A) Sequence alignments of sRNAs in the sR09 family is presented and their guide complementarities to 23S rRNA are indicated below. The critical “N+five” nucleotide in the guide regions is highlighted in blue when there is a Watson-Crick base pair between the guide and target and in rose when there is a mismatch base pair. Mismatched base pairs are also indicated by an asterisk in the rRNA position. The yellow highlight represents the “N+five” position in the rRNA target. (B) The secondary structure of helix 68-69 region in 23S rRNA is depicted showing the complementarity of the sR33 D guide near position G2021 and the Dʹ guide near position C2045. All six sR33 members have a mismatch at position 2045 at the “N+five” position in the guide-target interaction. Secondary structure generated by SSU-ALIGN package (55).

In three cases, only one member of a family has a mismatched base pair (*Pis* sR106:A608, *Par* sR09:U912, and *Pae* sR44:C2117, all in 23S). For example, the sR09 family has five members and the D and D′ guides are highly conserved, yet the *Par* sR09 D guide contains an A-to-U nucleotide substitution at the “N+five” position, changing the guide-target interaction to U:U at this position (Supplementary Figure S8). These guide-target interactions may still play an important role in the localized folding of the 23S rRNA in each species, even if one guide occassionally contains a mismatch impairing methylation (Supplementary Figure S8B).

#### Guides with no predicted targets in tRNA or rRNA (orphan guides)

In the 526 different *Pyrobaculum* sRNAs, 28% of guides (292/1052) show no significant complementarity to either rRNA or tRNA sequences (see Supplementary Tables S4 and S5). We also searched for mRNA and other non-coding RNA targets for these orphan guides, but no significant, conserved complementarities were observed, including orphan guides conserved across all six *Pyrobaculum* species.

In some instances, orphan guides appear to be the result of a small number of nucleotide substitutions. For example, in the sR30 family, the D guide of all six members is predicted to target C2724 in 23S rRNA, whereas the Dʹ guide is predicted to target C2708 in only four of the members (Supplementary Tables S1, S4). The two sRNAs containing the diverged Dʹ guides (in the *Pog* and *Par* sub-lineage) contain just two changes at the beginning of the guide region, dropping the guide-target pairing interaction to below the minimum threshold of eight base pairs. These “orphan guides” that are part of ancestral dual guide sRNAs may in fact still help mediate rRNA folding, albeit with a weaker interaction.

Guide divergence also appears to occur via genomic arrangements or sRNA duplication that result in an overlap between an sRNA gene and a protein-coding gene. Of the sRNAs that overlap the 5ʹ- or 3ʹ-end of a protein-coding gene in the sense orientation, 88% and 61% of the respective overlapping guides do not have predicted targets in rRNA or tRNA (see later sections for more discussion).

The origin of other orphan guides is less clear. There are a few instances where guide families are conserved but only one of the members has targets. For example, both *Pae* sR64 and *Pae* sR101 have tRNA targets, but the five other homologs in their families are dual orphan guides (Supplementary Tables S4 and S5). These guide families are relatively well conserved with only point mutations, and it is unclear if these are instances of gain- or loss-of-targeting function. There are only three families (sR43, sR50 and sR108) in which all six members have dual orphan guides. Interestingly for sR43, both guides are highly conserved among all members and so would be expected to recognize the same two targets, if they could be identified.

### Proliferation, mobility, plasticity and evolutionary divergence of C/D box sRNA genes within the *Pyrobaculum* genus

Grouping the *Pyrobaculum* C/D box sRNAs into 110 homologous families has greatly facilitated our ability to observe evolution of sequence features at a time-scale where variation is plentiful but homology can usually be established, even for mobilized sRNA genes. Here we describe sequence similarity of guide sequences, which impart function and are the most conserved, defining elements of homologus families.

#### Composite and transposed sRNAs

Of the 18 sRNA single-member families, five have a guide that shares some resemblance to a guide in a different sRNA family. These are designated as either transposed or composite sRNAs (Table 1). Transposed sRNAs share one guide with another defining family, but typically the guide has been transposed from D to Dʹ or visa versa, from Dʹ to D position, compared to the defining family. Composite sRNAs have D and Dʹ guides that each match one of the guides in two different families. For example, *Pca* sR12/45 has a D guide that is similar to the D guide of the sR12 family and a Dʹ guide that is similar to the Dʹ guide of the sR45 family (Figure 4). We suggest that the genes encoding these composite and transposed sRNAs are generated by genomic rearrangements between different sRNAs or sRNA genes.

**Figure 4:**
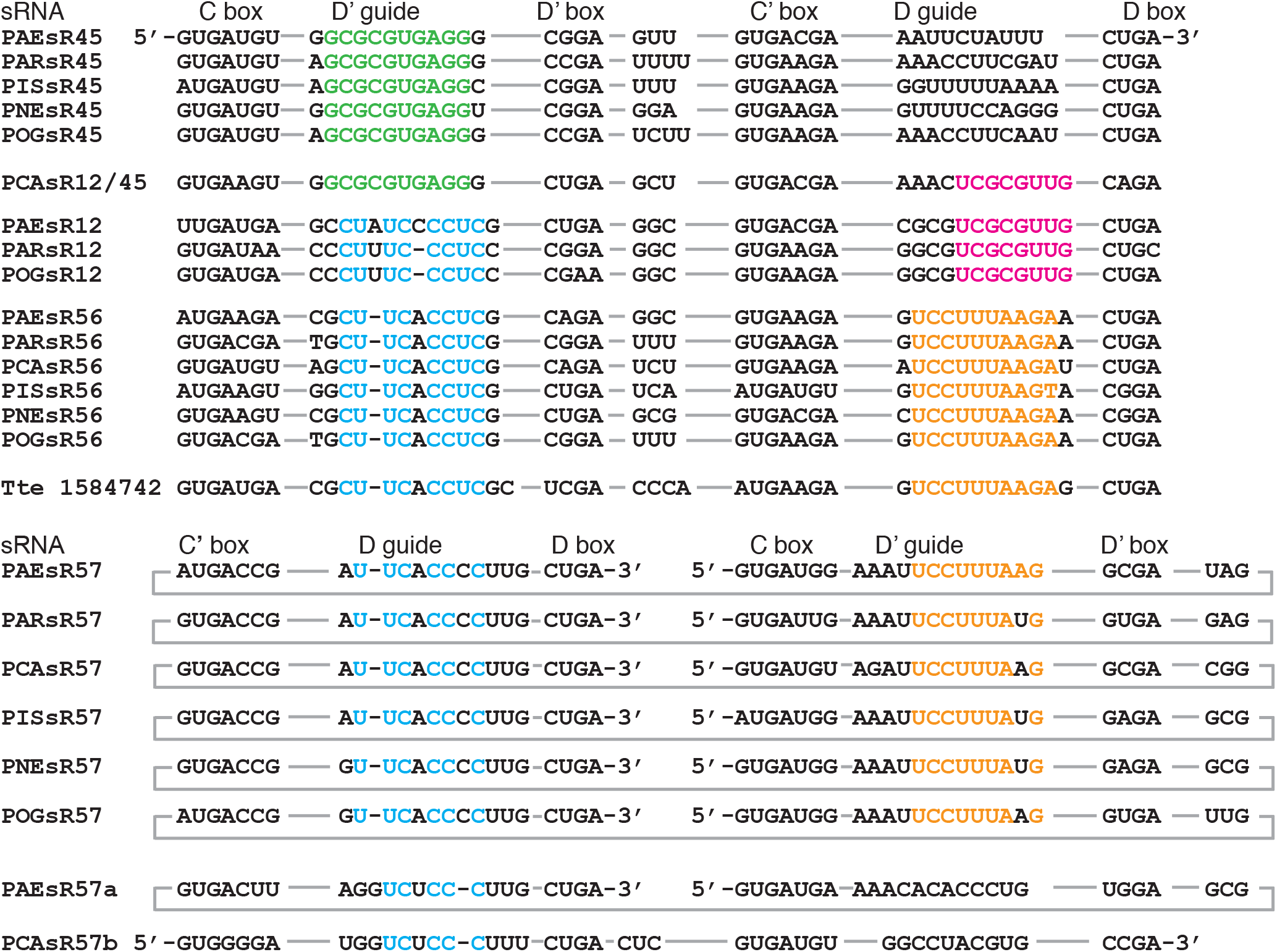
Interconnected guide sequence similarity between different families of sRNAs. The colored sequences (green, magenta, blue and orange) indicate different sequence similarities in the guide regions of sRNAs of the interconnected sRNA families sR45, sR12/45, sR56, and sR57. The sRNA in the outgroup species *Thermproteus tenax* (*Tte*; chromosome start 1584742) is related to the sR57 family.

**Table 1:** Composite and transposed C/D box sRNAs. Two unusual types of sRNAs (composite and transposed) were identified. Composite sRNAs have a D guide that shows sequence similarity to a guide in one sRNA family, whereas the D′ guide shows sequence similarity to a guide from a second sRNA families. These are given both family numbers separated by a forward slash (/). Transposed sRNAs have either a D guide that is shared with the D′ guide of the defining family, or visa versa, a D′ guide that is shared with the D guide of another family. Transposed sRNAs are identified with the number of the defining family followed by a lower-case a or b. The *Pae* sR57b is considered as a transposed sRNA since the D′ guide normally associated with the sR57 is not present (to view these sRNA sequences, see <u>http://lowelab.ucsc.edu/snoRNAdb/)</u>.

| Composite<br>C/D box sRNA | D' guide | D guide |
| --- | --- | --- |
| *Pca* sR12/45 | Shared with D' guide of sR45 family | Shared with D guide of sR12 family |
| *Pca* sR103/109 | Shared with D' guide of sR103 family | Shared with D guide of sR109 family |
| Transposed<br>C/D box sRNA(s) | D' guide | D guide |
| *Pca* sR26a | Shared with D guide of sR26 family | Not shared |
| *Pog* 46a | Shared with D' guide of sR46 family | Not shared |
| *Pca* sR57a | Shared with D guide of sR57 family | Not shared |
| *Pae* sR57b | Not shared | Shared with D guide of sR57 family |
| *Par* and *Pog* sR13a | Shared with D guide of sR13 family | Not shared |

#### Duplication of sRNAs

Duplication of a full-length sRNA gene can also occur as evidenced by the highly similar *Pae* sR113a and 113b (Supplementary Figure 9A), a duplication not found in other species. The 5ʹ-flanking regions in front of the two genes are unrelated but the 3ʹ-flanking regions have substantial similarity. Downstream of the sR113a gene, sR08 is encoded on the opposite strand, yet the sR113b gene contains what appears to be the remnant of the sRNA gene that has been obliterated by the ORF PAE3005. The sR113a and sR08 genes are convergently transcribed and separated by a 1 bp intergenic space, a highly unusual arrangement where transcription of one may interfere with the other. There are small RNAseq reads for each of these sRNAs (sR113a, sR113b, sR08). Other examples of apparent duplications include the sR46 family, where only *Pog* contains a nearly identical sR46a gene, and *Pae* sR62, where an apparent pseudogene can be found about 2.5Kbp away (Supplementary Figure 9B).

#### Superfamilies of sRNAs illustrate guide evolution

The sR45, 12, 56 and 57 families appear to share a complex lineage, based on analysis of their guide regions (Figure 4). The sR56 and sR57 families represent a likely ancient duplication appearing early within the *Pyrobaculum* lineage. Both families have representatives in all six *Pyrobaculum* species, and one family (sR57) is a circular permutation of the other (sR56). The closest out-group species for *Pyrobaculum*, *Thermoproteus tenax* (*Tte*), contains only a homolog to sR56 out of these four sRNA families. The D and Dʹ guides of sR56 target modifications to 16S U877 and G908 respectively and the D and Dʹ guides of sR57 target modifications to 16S G906 and A879 respectively. The shared core sequence between the D guide of sR57 and the Dʹ guide of sR56 is UUCACC and the shared core sequence between the D guide of sR56 and the Dʹ guide of sR57 is UCCUUUA. These cores sequences are offset by two nucleotides due to indels within the respective guides and this accounts for the two nt shift in target specificities. The two aberrant (transposed) members of the sR57 family (*Pae* sR57a and *Pca* sR57b) are circular permutations of each other and share only a single guide (UC-CC-CUU, dashes indicate indels) with the core sR57 family. Archaeal sRNAs are known to circularize (29, 41–43) and the sR57 may be a prime example of circularization and re-insertion into the genome.

The sR12 family is also implicated in this complex interconnection of families. It has a Dʹ guide that exhibits sequence similarity to the Dʹ guide of the sR56 family (CU-UC-CCUC). Indels in the sR12 Dʹ guide changes the target specificity to position 23S G1221. As mentioned above, the D guide in the sR12 family is shared with the D guide of the composite sR12/45. The second Dʹ guide of sR12/45 is derived from the sR45 family; interestingly, this guide is predicted to target methylation to position C34 in the anticodon loop of tRNA^Val^.

The relationships between these four related families illustrate several important aspects of sRNA gene evolutions including: (i) gene duplication; (ii) target migration (resulting from insertion/deletion) or divergence (resulting from nucleotide substitution) that alters or abolishes guide-target interactions; and (iii) rearrangements, including guide replacement and/or circular permutation.

#### MITE-like elements resembling sRNAs

Many of the families with only one sRNA member occur in *Pca* (Figure 1C). This species exhibits modular duplications and rearrangements between and within sRNA families as evidenced by the transposed and composite sRNAs described above (see Table 1). A careful analysis has also revealed the presence of a MITE-like element present in at least 15 copies within the *Pca* genome (Figure 5, Supplementary Table S6). MITEs are miniature inverted-repeat transposon elements that are characterized by a combination of terminal inverted-repeats and internal sequences too short to encode proteins. These elements are Class II transposons that occur in plants and other archaea (44).

**Figure 5:**
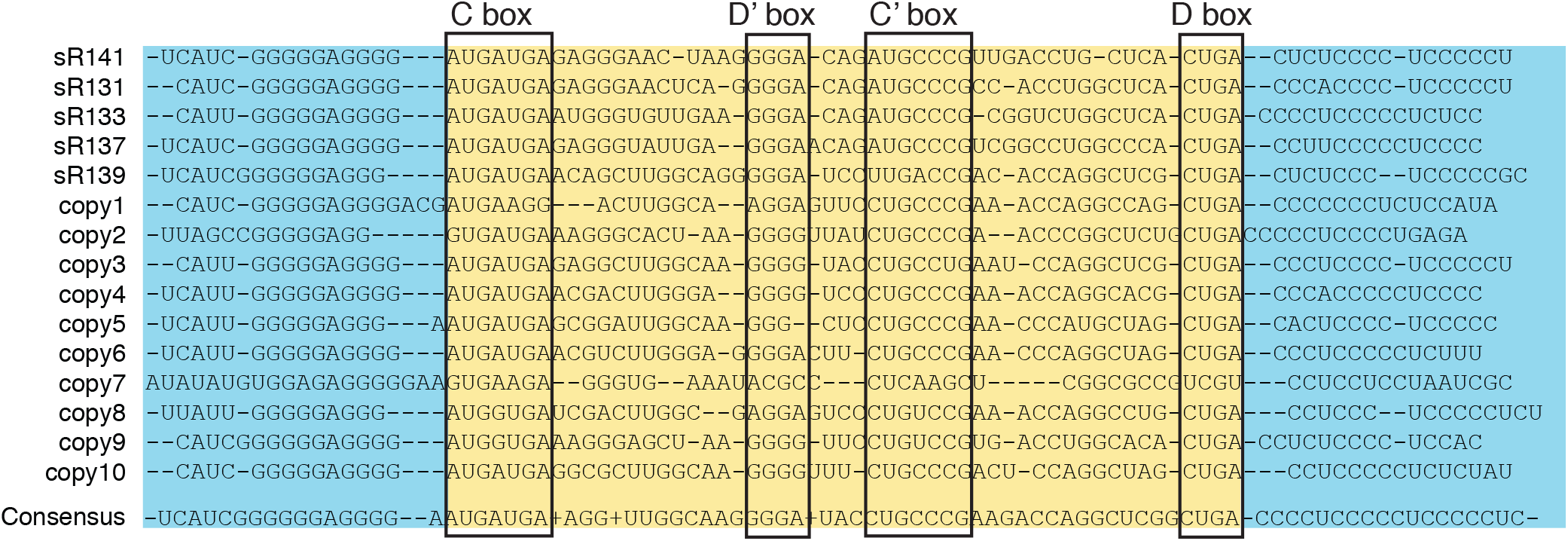
MITE-like element in the *Pca* genome. The chromosome of *Pca* contains fifteen copies of a MITE-like sRNA element. The sequences are aligned to illustrate the high degree of conserved sequence similarity in the 5ʹ and 3ʹ flanking inverted repeat sequences (blue highlight). The sRNA-like sequences (yellow highlight) contain canonical C and D boxes but generally degenerate Dʹ and Cʹ boxes (boxed). The conservation between the D and Dʹ guide sequences in the 15 elements is moderate with a consensus sequence at the bottom. Five of these elements were cataloged as authentic sRNAs.

In *Pca*, these elements have characteristics of both sRNAs and MITEs. Each identified copy contains near-consensus C box and D box sequences, but highly degenerate internal Dʹ and Cʹ sequences. Both guide sequences (adjacent to D and D’ boxes) exhibit only modest sequence similarity across the 15 copies. Highly conserved imperfect inverted-repeat sequences flank the C and D boxes (Figure 5). The elements have a large average distance (322 nt) from the nearest protein-coding gene compared to other sRNAs (22 nt). The presence of these MITE-like elements in regions of the genome where there are no other annotated genomic features *and* that do not align with other *Pyrobaculum* genomes suggests that they are located in regions of genomic instability, natural hotspots for insertion by mobile elements.

Five of the element copies were classified as C/D box sRNAs (sR131, sR133, sR137, sR139, sR141) and contain moderately degenerate internal box sequences. The genomic locations of another ten copies of this element, which were too diverged to be considered as potential sRNA genes, are listed in Supplementary Table S6. A phenomenal 13.6% of the uniquely mapped RNA-seq reads in *Pca* were generated from a single MITE-like locus (sR141). For the other *Pca* sRNAs (non-MITE-like), the highest percentage of uniquely mapped reads to an sRNA was 4%, and on average 0.5% of total unique reads mapped to each sRNA. The other MITE-like elements have abundance levels similar to other sRNAs. The MITE-like sRNAs also tend to have a higher percentage of antisense reads compared to other sRNAs. On average, 39% of reads from a MITE-like sRNA locus are antisense, whereas on average 9.3% of reads from other *Pca* sRNAs are antisense. We suggest that this element may play a role in the generation, mobilization and proliferation of C/D box sRNAs or their modular components. We did not detect these MITE-like elements in the other five *Pyrobaculum* species, although they may exist in lower copy numbers and with greater sequence divergence.

#### Association of C/D box sRNAs with other non-coding RNAs

Most archaeal C/D box sRNAs are independently transcribed, but in a few cases C/D box sRNA genes are known to be polycistronic (1). Transcription of archaeal C/D box sRNAs genes with protein-coding genes has been reported in *Sulfolobus* and *Pyrococcus* species (16, 19); in *Nanoarchaeum equitans*, a few instances of di-cistronic C/D box sRNA-tRNA transcripts have also been reported (45).

Among the different *Pyrobaculum* species, the transcriptional relationships between sRNA and other non-coding RNA genes can be dynamic. We identified a novel, three gene C/D box sRNA polycistron (sR101, sR21, and sR100) in *Pae,* confirmed by northern blot (Supplementary Figure S10), which appears to be shared in *Pis* and *Pca* based on genomic proximity and overlapping RNA-seq reads (Figure 6A). In contrast, three species (*Pne*, *Par*, *Pog*) have lost sR100, but maintain synteny between sR21 and sR101. In *Pis* and *Pne* the sR34 and sR40 genes are also co-transcribed (based on RNA-seq reads) and separated by 10, and -4 nts respectively. In *Pne* the D box of sR34 is located within the C box of sR40 (4 nt overlap); it is unclear how this overlap affects the maturation of the two sRNAs. In *Par*, *Pog*, and *Pca* the genes are separated by 16, 16, and 78 nts respectively and are convergently transcribed whereas in *Pae* the two genes are separated by more than 2000 nts.

**Figure 6:**
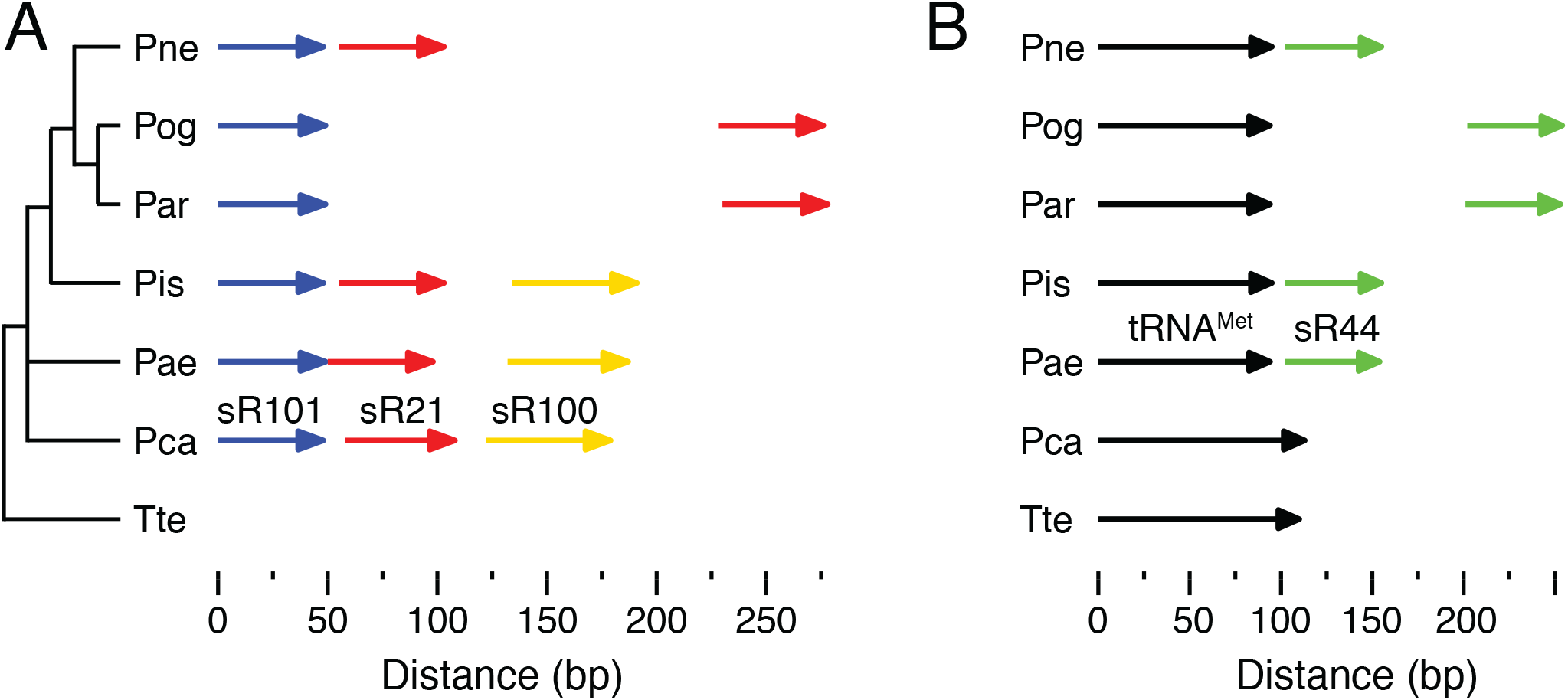
Genomic context of sRNA genes. (A) Genomic organization of the sR101 (blue), sR21 (red) and sR100 (yellow) genes in the six species of *Pyrobaculum*. The 16S rRNA phylogenetic tree with *Thermoproteus tenax* (*Tte*) as the outgroup, is illustrated on the left; the sRNA gene locations above a bp distance scale is illustrated to the right for the *Pyrobaculum* species. There is no representative of the sR100 gene in *Pne*, *Pog*, *Par* and in *Par* and *Pog* there is an approximately 200 nt insertion between the sR101 and sR21 genes. (B) Linkage of tRNA and sRNA genes. In *Pae*, *Pis*, and *Pne* the sR44 genes (grey arrows) are located eight nts or less from the 3ʹ end of a tRNA^Met^ gene (green arrows). In *Pog* and *Par* the distance between the tRNA^Met^ and sR44 gene is increased to about 100 nts and appear to be expressed from separate promoters. In *Pca*, sR44 is located approximately 13 Kbps downstream of the tRNA^Met^ gene. There is no representative of the sR44 family in *Tte*.

Plant species and the archaeon *Nanoarchaeum equitans* have C/D box sRNA genes that are reported to be co-transcribed with tRNAs (45). In the *Pyrobaculum* genus, we find a single case of a C/D box sRNA family (sR44) that is likely co-transcribed with elongator tRNA^Met^, although the spacing between the two genes is extremely variable, ranging from 8 nucleotides (*Pae, Pis, Pne*) apart to 100 nt (*Par, Pog*) to 13 Kbp in *Pca* (Figure 6B).

These examples demonstrate fluidity of C/D box sRNA co-transcription with other non-coding RNAs within the *Pyrobaculum* genus and preference for individual promoters. None of the four sRNAs discussed (sR21, sR100, sR101, and sR44) have homologs in the out-group species *Tte*, so these sRNAs probably arose in the *Pyrobaculum* lineage (Figure 6). Within the polycistronic example, the sR100 was lost from the transcription unit in the *Par*/*Pog*/*Pne* lineage and in *Pog* and *Par* the remaining sR100 and sR101 genes developed individual promoters. Similarly, the sR44 gene appears to have become linked to the tRNA^Met^ gene in the ancestor of *Pae*, *Pis*, *Pne*, *Pog*, and *Par* lineage; *Pca* is an out group to these species and does not have the same sRNA-tRNA linkage (Figure 6B). Separate promoters for the two genes occur in the *Pog*/*Par* sub-lineage.

### Impact of overlapping of C/D box sRNA genes and protein-coding genes

In a previous study (18), we noted that *Pyrobaculum* C/D box sRNAs genes are over 40-fold more likely than tRNA genes to have conserved overlap with orthologous protein-coding genes. Other studies have also noted the 3ʹ-antisense overlap of C/D box sRNAs with protein-coding genes (16). We looked more closely at this relationship because overlap could impact the function of both gene types. In addition, antisense interactions suggest the possibility that C/D box sRNAs might guide modification of mRNAs or be involved in antisense regulation.

In the set of 526 *Pyrobaculum* C/D box sRNAs, 97 exhibit either partial or complete overlap with protein-coding genes (Figure 7, Supplementary Figure S11 and Supplementary Table S7). For this analysis, we considered only overlaps that extend either into the Dʹ guide region (eight nts or more beyond the 5ʹ-end of the sRNA gene) or into the D guide region (five nts or more beyond the 3ʹ-end of the sRNA gene) since shorter overlaps ending in the D box or C box were not expected to impact target specificity. We note, however, there are many instances (23 total) where the 5’ end of a slightly overlapping downstream sRNA gene (same strand) provides the translation stop codon for the upstream ORF (e.g., C box R<u>UGA</u>UGA). We classified the more extensive overlaps into seven categories (Figure 7, Supplementary Figure 11 and Supplementary Table S7). Instances of overlap with the 5ʹ-end of an mRNA were checked manually to confirm that the start codon of the mRNA was called correctly; start codons were adjusted based on protein sequence conservation.

**Figure 7:**
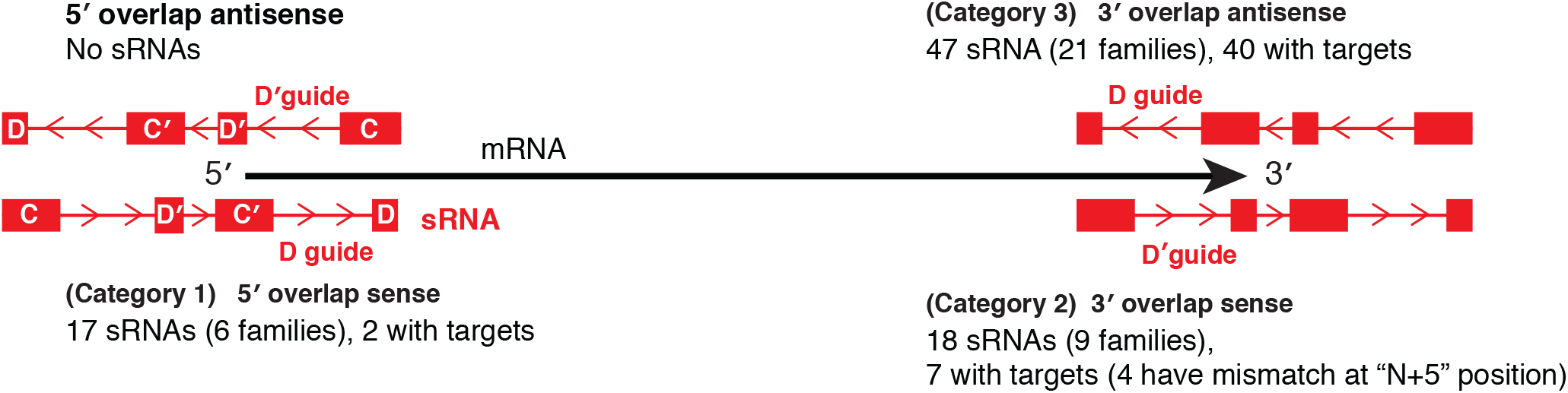
The overlap between sRNA genes and protein coding genes. The overlap between sRNA genes and protein-coding genes is divided into seven categories. The first three are shown here and the rest are in Supplementary Figure S11. The protein genes are shown as black arrows with the 5ʹ and 3ʹ polarity indicated. Overlapping sRNA genes are shown in red with polarity indicated by the internal arrows; the C, Dʹ, Cʹ and D box sequences are indicated as shown in the top left sRNA. The number of sRNA genes, the number of families that they represent and the number that have predicted targets is indicated for each type of overlap. Details relating to these sRNAs are given in the text and in the Supplementary Table S7.

The major three categories of sRNA genes that overlap a protein-coding gene involve 82 genes: (category 1) overlap at the 5ʹ-end of the protein ORF in the sense orientation; (category 2) overlap at the 3ʹ-end of the protein ORF in the sense orientation and (category 3) overlap at the 3ʹ-end of the protein ORF in the antisense orientation (Figure 7). There were no C/D box sRNA genes that overlapped the 5ʹ-end of a protein-coding gene in the antisense orientation. In category 1, only two of the 17 overlapping guides (12%) were predicted to have methylation targets in rRNA or tRNA. We suspect that in many of these instances, the sRNAs are co-transcribed with the mRNA based on promoter analysis. The translation initiation codons for the respective ORFs are located either in the Cʹ box or in the D guide region of the sRNA sequence. A recent study by Tripp *et al*. reached a similar conclusion based on an analysis of 300 sRNAs from six divergent species of archaea (46).

The second and third categories with overlapping guides in the sRNAs at the 3ʹ-end of the protein-coding gene had, in comparison, numerous predicted targets (40 of 47 for antisense sRNAs guides and 7 of 18 for sense sRNA guides; see Supplementary Table S7). This disparity suggests the 3ʹ-end of protein-coding genes is more flexible and better accommodates both amino acid sequence encoding and sRNA guide function.

The high proportion of sRNAs located near or overlapping the 3ʹ-end of protein-coding genes may suggest that they play a role in gene regulation and possibly mRNA stability. Sense strand sRNAs that are co-transcribed with mRNA need to be excised and rescued from decaying mRNA transcripts. The sRNAs that are antisense could participate in antisense regulation through the formation of an RNA/RNA duplex or trigger methylation of the mRNA through a more limited guide target interaction. We also note in the RNA-seq reads that many sRNA genes generate both sense strand and antisense strand transcripts. In other archaea, small antisense RNAs have been shown to regulate gene expression by binding to 3ʹ-UTRs (reviewed in (47)). A role for these antisense sRNA transcripts has not been defined, in part because there is no experimental genetic system for *Pyrobaculum*.

The remaining sRNA-mRNA overlap categories are much less common, collectively involving 15 genes. Category 4 represents sRNAs that are contained completely within protein-coding genes (Supplementary Figure S11A). Nine of the ten of these are in the antisense category and all have at least one guide that has a target in rRNA or tRNA. These internal sRNAs are located near the 3ʹ-end of the protein-coding gene, again suggesting that this region is flexible and can accommodate both amino acid coding and guide function without detriment. Categories 5-7 include sRNA genes that overlap two adjacent mRNA-encoding genes (Supplementary Figure S11B-D). These types of overlaps are rare and only 1-2 sRNAs fall into these categories (see Supplementary Figure S11 for more discussion).

In summary, our analyses indicate that overlap of an sRNA gene at the 5ʹ end of an ORF in the sense orientation is generally not compatible with the targeting function of the overlapping guide, but overlap on the 3ʹ end of an ORF in either sense or antisense orientation is much more common. Some families such as sR05, sR118, and sR3 have conserved overlap (Supplementary Table S7). However, there are many more instances where only a subset of the sRNAs in a family have conserved overlap, indicating that the position of sRNAs in relation to ORFs can be dynamic. Some orphan guides may be a result of loss-of-function by overlap with an ORF, rather than the result of developing targets other than tRNA or rRNA.

## DISCUSSION

In the last five years, the scope and number of archaeal C/D box sRNAs has been more fully realized with high-throughput RNA-seq data (17, 18). We took advantage of this data, along with computational methods and comparative genomics, to identify a comprehensive set of 526 C/D box sRNAs from six species within the genus *Pyrobaculum*. We organized these sRNAs into 110 homologous families based on sequence similarity and predicted their methylation target sites across both rRNA and tRNAs. With this set of families and predicted targets, we were able to explore known and hypothetical functions of C/D box sRNAs, study their impact on genomic organization and architecture, and visualize many aspects of their evolutionary origins and diversification. We have identified instances of C/D box sRNA gene expansion and diversification resulting from a number of processes: (i) gene duplications, (ii) gene rearrangements, (iii) guide replacement or translocations, (iv) transposon-like mechanisms, and (v) guide divergence caused by nucleotide mutations, insertions or deletions.

With our curated set, we developed a new computational model for detection of archaeal C/D box sRNAs. This collection provided a more comprehensive training set than what was available for prior computational search methods (22), and allowed us to incorporate K-motif information. We were able to detect new C/D box sRNAs in *Pyrococcus* species, well-studied organisms from a different archaeal phylum, indicating that this model and using expanded training data should be an effective strategy for detection of C/D box sRNAs in other archaeal phyla, the focus of follow-up studies.

The combination of the *Pyrobaculum* C/D box sRNA catalogue and our extensive map of the corresponding methylation sites provides evolutionary perspective on the canonical functions of archaeal C/D box sRNAs. Our analysis indicates that slightly less than two-thirds of the predicted targets are conserved among the six species. This is in contrast to target conservation as described in a recent panarchaeal study where only one target was conserved among species from seven different orders (17). This study also shows that at the genus level, a core set of methylation sites are highly conserved, but a consistent 10-20% continue to change relative to other species. There are several instances of conserved *Pyrobaculum* sRNA families where members have slightly different targets because of insertions or deletions within one of their guides (e.g., sR127; Supplementary Figures S1, S2 and Supplementary Table S4). In these and other instances, the region of interaction within rRNA is conserved but the particular site of methylation is not. There are intriguing instances of dual guide target long-range interactions, but these are also rarely conserved past the genus level. We imagine that these interactions help organize localized regions within the tertiary 16S or 23S rRNA structures. The variation in methylation sites is reflective of the sequence evolution of the guides and reinforces the hypothesis that the aggregate of methylations in certain regions of the rRNA is generally more important than particular sites of modification (9).

Examination of sRNA genomic context and contrasting those targeting tRNAs versus rRNAs provides new perspective on why some guides do not appear to target modifications. Many C/D box sRNAs that target rRNA are dual guides (260/327, 80%) and for most of these (237/327, 72%), both targets are within 100nt of each other on rRNA, strongly implicating them in ribosome assembly. In contrast, only about 20% of sRNAs that target tRNAs (29/144) are dual guides. This suggests that a dual guide sRNA is generally not advantageous for guiding methylation of tRNAs. Another case where dual guides are not common is when an sRNA overlaps with a protein-coding gene or a promoter. In this case, the protein-coding function of overlapping sequence is more strongly selected than C/D box sRNA guide function, leaving only the non-overlapping guide region to target tRNA or rRNA. There are also a few instances in our dataset where there are dual orphan guides, so it is possible that these sRNAs serve another purpose other than guiding methylation of tRNA or rRNA.

The extensive overlap of C/D box sRNAs genes with protein-coding genes raises new questions about their sequence constraints, excision from mRNA transcripts, and role in the regulation of mRNA stability and translation. In numerous instances, the modification function of sRNA guides is overridden by the coding constraints of the mRNA sequence, particularly at the 5ʹ end of overlapping protein-coding genes. Sense strand sRNAs are interesting because their maturation requires precise excision from the mRNA transcript, and could conceivably create an alternate start codon downstream. Alternatively, if the sRNA sequence is not excised from the mRNA, it could interfere with translation initiation by attracting L7Ae binding, and possibly also Nop 56/58 and fibrillarin assembling on the mRNA. The RNP complex likely protects the sRNA sequence from nucleases that degrade the unprotected parts of the mRNA transcript, allowing precise excision of the sRNA from the mRNA transcript. Tripp *et al.* have recently made a similar suggestion, and using artificial constructions, provided experimental evidence to support this idea (46). Preliminary data from our lab also supports the hypothesis that K-turns form in archaeal mRNAs and are bound by L7Ae (27). The K-turn is known to regulate mRNA translation by protein-binding in both natural and synthetic constructions (48–50). In contrast to sense sRNAs, antisense sRNAs are intriguing when they overlap the 3ʹ end of an mRNA. These antisense sRNAs may function as antisense regulators, or may use their guide sequences to carry out site-specific methylation of the mRNA and influence the structure, function or stability of the mRNA.

This comprehensive effort to identify the complete set of C/D box sRNA genes from six species within the hyperthermophilic genus *Pyrobaculum* has provided unique and valuable insights into (i) their structure and function, (ii) their role in ribosome subunit biogenesis, (iii) their evolutionary persistence, propagation and divergence and (iv) their potential roles in influencing protein gene expression and shaping overall genome architecture.

## ACKNOWLEDGEMENTS

The authors thank Dr. Patricia Chan for her helpful comments with the manuscript and assistance with uploading data to the GEO database.

